# In search of the RNA world on Mars

**DOI:** 10.1101/2020.02.28.964486

**Authors:** Angel Mojarro, Lin Jin, Jack W. Szostak, James W. Head, Maria T. Zuber

## Abstract

Advances in origins of life research and prebiotic chemistry suggest that life as we know it may have emerged from an earlier RNA World. However, it has been difficult to reconcile the conditions used in laboratory experiments with real-world geochemical environments that may have existed on the early Earth and hosted the origin(s) of life. This challenge is in part due to geologic resurfacing and recycling that have erased the overwhelming majority of the Earth’s prebiotic history. We therefore propose that Mars, a planet frozen in time, comprised of many surfaces that have remained relatively unchanged since their formation >4 Gya, is the best alternative to search for environments consistent with geochemical requirements imposed by the RNA world. In this study we synthesize *in situ* and orbital observations of Mars and modeling of its early atmosphere into solutions containing a range of pHs and concentrations of prebiotically relevant metals (Fe^2+^, Mg^2+^, and Mn^2+^), spanning various candidate aqueous environments. We then experimentally determine RNA degradation kinetics due to metal-catalyzed hydrolysis and evaluate whether early Mars could have been permissive towards the accumulation of long-lived RNA polymers. Our results indicate that a Mg^2+^-rich basalt sourcing metals to a slightly acidic (pH 5.4) aqueous environment mediates the slowest rates of metal-catalyzed RNA hydrolysis, though geologic evidence and modeling of basalt weathering suggest that aquifers on Mars would be near neutral (pH ∼7). Moreover, oxidizing conditions on Mars have major consequences regarding the availability oxygen-sensitive prebiotic metals (i.e., Fe^2+^ and Mn^2+^) very early in its history due to increased RNA degradation rates and precipitation. Overall, 1) low pH better preserves RNA than basic conditions at high concentrations; 2) acidic to neutral pH environments with Fe^2+^ or Mn^2+^ will hydrolyze more RNA; and 3) alkaline environments with Mg^2+^ dramatically hydrolyze more RNA.

## 1. Introduction

The origins of life can best be understood as a series of plausible steps culminating in the emergence of a “self-sustaining chemical system capable of Darwinian evolution” (NASA Astrobiology). However, life as we know it is a highly complex collection of molecular machinery and genetic information. The central dogma of molecular biology stipulates that deoxyribonucleic acid (DNA) makes ribonucleic acid (RNA) via transcription, RNA makes protein via translation, and information cannot be transferred backwards from proteins to nucleic acids (Crick, 1970). Fundamentally, neither DNA, RNA, nor proteins can exist without the others as they do today. Nevertheless, this dilemma belies the fact that the capability of translating information between dissimilar polymers (e.g., polynucleotides to polypeptides) is mediated by the ribosome, an RNA enzyme (Cech, 2000). This is significant because the ribosome is arguably an evolutionary anachronism from a period where RNA polymers acted as both enzymes (protein) and information storage (DNA) (Petrov et al., 2015). Additional discoveries of structural and regulatory RNA molecules (Breaker, 2012) suggest that life may have emerged from an earlier RNA world dominated by ribozymes (e.g., the ribosome) (Gilbert, 1986) and ribonucleotide-containing molecules (e.g., adenosine triphosphate - ATP) (Hernández-Morales et al., 2019) catalyzing reactions and mediating a protometabolism.

Under the RNA world scenario, abiotic synthesis of simple RNA molecules from common molecular feedstocks in geologically relevant environments (e.g., Patel et al., 2015) would have given rise to self-assembling protocellular systems (Joyce and Szostak, 2018). Thereafter increasingly complex RNA polymers capable of both hereditary storage and autocatalysis would precede the DNA-RNA-protein world (Bernhardt, 2012). To test the RNA world hypothesis investigators have experimentally demonstrated: 1) abiotic RNA synthesis (e.g., Powner et al., 2009); 2) non-enzymatic RNA replication (Adamala and Szostak, 2013; Jin et al., 2018); 3) self-assembly of protocell membranes and membrane replication (Schrum et al., 2010); 4) directed evolution and fitness landscapes yielding persistent RNA motifs (Jimenez et al., 2013) and functional ribozymes (Voytek and Joyce, 2007); and 5) the co-synthesis of RNA, amino acids, and lipids (Patel et al., 2015).

Although strides in prebiotic chemistry have demonstrated the viability of an origin of life via the RNA world, a longstanding criticism is that RNA is inherently unstable due to the presence of a nucleophilic 2’-hydroxyl group which readily catalyzes cleavage of the 5’,3’-phosphodiester bond (Li and Breaker, 1999). Because of this characteristic, RNA is deemed an ephemeral molecule that is unlikely to accumulate, functionalize, and precipitate life in a prebiotic world. Researchers have therefore directed efforts towards determining particular conditions or cofactors which can stabilize RNA in real-world environments. Experimental work suggests: 1) RNA is most chemically stable between pH 4 – 5 (Bernhardt and Tate, 2012; Oivanen et al., 1998) and near 0 °C (Kua and Bada, 2011; Levy and Miller, 1998); 2) metal cofactors such as Fe^2+^, Mg^2+^, and Mn^2+^ facilitate the folding of RNA polymers into stable secondary and tertiary structures (Bowman et al., 2012; Laing et al., 1994; Petrov et al., 2012); 3) copolymers such as polypeptides and polysaccharides can favor specific polynucleotide conformations, resulting in persistent structures and vice-versa (Runnels et al., 2018); and 4) folding of many RNA sequences decreases rapidly above 30 °C (Moulton et al., 2000).

Still, it has been difficult to confidently determine the dynamic environments that could have existed on the Hadean Earth and hosted the origin of life. This challenge is in part due to geologic resurfacing and recycling that have erased the overwhelming majority of the Earth’s prebiotic history (Marchi et al., 2014). Nevertheless, we can speculate that the likeliest time interval for the origin of life on Earth is best constrained by the accretion of the first continents following a Moon-forming impact 4.5 – 4.3 Gya (Monteux et al., 2016) and the earliest putative biosignatures ∼3.7 Gya (Nutman et al., 2016). Given this consideration, the best alternative is to search for environments consistent with RNA stability on Mars, a planet frozen in time, preserving primordial surfaces which have remained relatively unchanged since they formed >4 Gya (Hartmann and Neukum, 2001). Reminiscent of the Hadean epoch on Earth (Nisbet and Sleep, 2001), the Noachian period on Mars is characterized by meteoritic bombardment and punctuated aqueous activity resulting in extensive ground water circulation (Ehlmann et al., 2011), valley networks (Fassett and Head, 2011), and long-lived lacustrine environments (Goudge et al., 2015, 2012; Grotzinger et al., 2014). Some researchers would argue that life began on Mars and was transported to the Earth around the timing of the earliest biosignatures ∼3.7 Gya (Benner and Kim, 2015). That is, non-sterilizing lithological exchange between Mars and Earth from impact ejecta produced during the presumed Late-Heavy Bombardment period (Boehnke and Harrison, 2016; Gomes et al., 2005) may have transported viable microbes between planets resulting in ancestrally related life (Gladman et al., 1996; Weiss, 2000).

The case for an origin of life on Mars relies on prebiotic environments that are inferred to be analogous to environments on Earth, common molecular feedstocks (including cometary sources) (Callahan et al., 2011), and plausible reactive pathways predicted on Earth that are applicable on Mars (e.g., Ritson et al., 2018) which may have resulted in parallel events in accordance to the RNA world hypothesis (Benner and Kim, 2015). This notion is further supported by *in situ* detection of boron (Gasda et al., 2017), which is considered crucial to stabilize ribose in the formose reaction (Furukawa and Kakegawa, 2017), experimental work that predicts higher phosphate bioavailability on Mars (Adcock et al., 2013), and the detection of clays (Ehlmann et al., 2011) that have been demonstrated to assist in non-enzymatic RNA polymerization (Ferris, 2006). This Mars origin of life hypothesis suggests that past or present Martian life may have utilized known building blocks (e.g., nucleic acids, sugars, amino acids) and closely resembled life as we know it. Moreover, if life exists on Mars today, it could theoretically be detected by means of nucleic acid (DNA and RNA) sequencing (Carr et al., 2017; Mojarro et al., 2019). Assuming that viable RNA was being delivered to Mars via unspecified sources (e.g., cometary or *in situ* synthesis) to UV-shielded aqueous environments (Cockell et al., 2000), here we investigate whether early Mars was permissive towards the accumulation of long-lived RNA polymers. We anticipate our findings could provide insight into potential mechanisms, environments, and requirements necessary for sustaining an RNA world on the early Earth.

## 2. Materials and Methods

### 2.1. Approach

The surface of Mars displays evidence for alternating climate regimes at regional-to-global magnitudes that have evolved on variable time scales not dissimilar to the Earth (McLennan et al., 2019). In general, early Mars contained a broad range of geochemical environments (e.g., acidic to alkaline) primarily influenced by redox chemistry. In this study, we synthesize *in situ* and orbital observations and modeling of the early Martian atmosphere in order to extrapolate representative solutions containing a range of pHs and metals analogous to various candidate aqueous environments on Mars. Below we detail our experimental design, which involves incubating an RNA-DNA chimeric oligomer (simply referred to as the RNA oligomer) to quantify the hydrolysis rate of the 5’,3’-phosphodiester bond at a single ribonucleotide within the aforementioned solutions. The goal of this study is to understand the influence of bedrock composition (e.g., mafic-ultramafic, iron-rich, magnesium-rich, etc.) and subsequent weathering of prebiotically relevant metals (Fe^2+^, Mg^2+^, and Mn^2+^) which have been demonstrated to catalyze hydrolysis (Fedor, 2002), folding (Laing et al., 1994), and translation (Bray et al., 2018) on RNA stability. Furthermore, pH is simultaneously adjusted to reflect the composition of a hypothetical anoxic and CO_2_-dominated atmosphere at variable pressures in equilibrium with surface waters, an average acid vent, and alkaline vent (Kua and Bada, 2011). The end result is an analysis of single-stranded RNA stability and degradation kinetics in an array of simulated prebiotic geochemical spaces on Mars.

### 2.2. RNA oligomer

A hybrid RNA-DNA oligomer, 5’-Cy3-TTT-TTT-rCTT-TTT-TTT-3’, was designed to contain one ribonucleotide (r) in between deoxyribonucleotides, allowing us to quantify hydrolysis at a single cleavage site (Figure 1) (Adamala and Szostak, 2013). RNAse free and HPLC-purified oligos were ordered from Integrated DNA Technologies (IDT).

**Figure 1:**
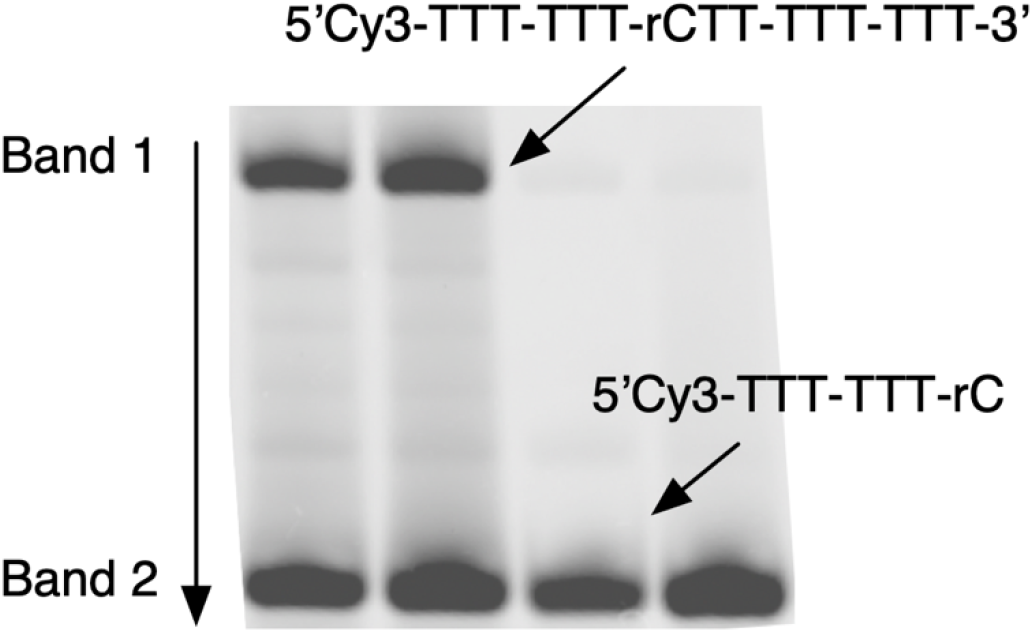
Oligonucleotide cleavage assay. A hybrid RNA-DNA oligomer, 5’-Cy3-TTT-TTT-rCTT-T TT-TTT-3’, was designed to contain a single ribonucletide (r) in between a chain of deoxyribonucleotides which could allow us to quantify cleavage at a single site. Representative gel scan displays two bands belonging to either the intact 15-mer (Band 1) or the residual 7-mer (Band 2) cleaved at the single ribonucleotide site.

### 2.3. Relevant observations

#### Mars Exploration Rover Opportunity

Sedimentary rocks exposed in the Meridiani Planum region (Burns formation) of Mars record alternating periods of acidic groundwater flow (pH ∼2 - 4) and desiccation under highly oxidizing conditions (Klingelhofer, 2004; McLennan, 2012; Squyres et al., 2006; Squyres and Knoll, 2005).

#### Mars Exploration Rover Spirit

Widespread Fe^3+^-sulfate soils (e.g., jarosite) at Gusev Crater indicate acid-weathering of primarily olivine-rich outcrops in a possible hydrothermal environment (Ming et al., 2008; Yen et al., 2008).

#### Mars Science Laboratory (MSL) Curiosity

Sedimentary rocks analyzed at Gale Crater (Bradbury-Mount Sharp groups) reflect a long-lived lacustrine environment with circumneutral pH waters and variable redox states as indicated by the presence of manganese oxide (Mn^2+^) deposits, magnetite-silica facies (Fe^2+^), and hematite-phyllosilicate facies (Fe^3+^) (Grotzinger et al., 2015, 2014; Hurowitz et al., 2017; Lanza et al., 2016).

#### Mars Reconnaissance Orbiter & Mars Express

Crustal Fe-Mg smectites indicate global groundwater circulation (Ehlmann et al., 2011) consistent with recent work suggesting subsurface waters were primarily anoxic, Fe^2+^-rich, and circumneutral pH which became rapidly acidified due to atmospheric O_2_ or photo-oxidation of Fe^2+^ to Fe^3+^ at the Martian surface (Hurowitz et al., 2010). Mg^2+^-rich carbonate deposits near Nili Fossae indicate neutral to alkaline pH waters likely in contact with a CO_2_ atmosphere (Ehlmann et al., 2008). Abundant Late Noachian aqueous surface environments and features, including fluvial valley networks (e.g., Fassett and Head, 2008; Hynek et al., 2010) open-basin lakes (e.g., Fassett and Head, 2008b; Goudge et al., 2012) and closed-basin (endorheic) lakes (e.g., Goudge et al., 2015).

#### Mars Atmosphere and Volatile Evolution

Isotopic evidence indicates a continuous loss of a ≥0.5 bar CO_2_-dominated and increasingly oxidizing atmosphere (Jakosky et al., 2017) since the early Noachian ∼4.1 Gya due to erosion by solar wind when the Martian dynamo is thought to have shut down (Lillis et al., 2013).

#### Atmospheric Modeling

No atmospheric model has been able to resolve liquid water on the early Mars with a faint young sun. Various models have suggested a range of atmospheric compositions (e.g., H_2_, CO_2_, H_2_O, SO_2_, H_2_S); however, none has been able to balance counteracting cooling effects of atmospheric density and albedo due to aerosol and cloud formation (e.g., Palumbo et al., 2018; Tian et al., 2010). Due to lower solar luminosity values in early history, recent atmospheric general circulation models (e.g., Forget et al., 2013; Palumbo and Head, 2018; Wordsworth et al., 2013) have found it difficult to maintain the >273 K mean annual temperature (MAT) seemingly required to support a “warm and wet” or “warm and arid” early Mars climate proposed to account for the valley networks interpreted to have transported liquid water on the surface (e.g., Craddock et al., 2003). Instead, these models suggest a “cold and icy” early Mars climate (Head and Marchant, 2014) in which snow and ice were deposited in the uplands, and episodic transient heating events caused melting and runoff to form the valley networks and lakes. Among the candidate transient events proposed are those due to: 1) spin-axis/orbital variations influencing peak annual and seasonal temperatures (Palumbo et al., 2018); 2) volcanic eruptions (e.g., Halevy and Head III, 2014); 3) impact events (e.g., Palumbo and Head, 2018; Segura et al., 2008; Steakley et al., 2019; Turbet et al., 2020); 4) subsurface radiolytic H_2_ production and release (e.g., Tarnas et al., 2018); and 5) collision-induced absorption (CIA) temperature amplifications during transient CO_2_ and methane release events (e.g., Wordsworth et al., 2017).

### 2.4. Mars prebiotic geochemical solutions

Given the aforementioned observations, we designed our experiments to simulate environments that are in equilibrium with an anoxic CO_2_-dominated atmosphere at variable pressures (10 bar, 0.1 bar, >>0.1 bar) inducing their respective shifts in pH (5.4, 6.7, 8) as calculated by Kua and Bada, 2011. Solutions representing an average acidic vent at pH 3.2 and alkaline vent at pH 9 were also included. Each pH solution contained 0, 2.5, 5, 10, 25, and 50 mM of Fe^2+^, Mg^2+^, or Mn^2+^ intended to represent a range of dissolved metal concentrations within the water column, variable weathering, and residence times. Mixtures of Fe^2+^ and Mg^2+^ at 50:50, 20:80, and 80:20 (25 mM total) were included to represent metal concentrations derived from variable bedrock compositions, in particular, 20:80 Fe^2+^:Mg^2+^ is closest to the average crustal composition on Mars (Mittlefehldt, 1994). A total of 5 pH conditions, 3 metals at 5 concentrations, 3 basalt analogs, and 5 negative controls resulted in 95 unique conditions.

Samples were prepared by mixing stock buffer solutions of 1 M Glycine-HCL (Sigma-Aldrich, 50046) at pH 3.2, 0.5 M MES (Sigma-Aldrich, 76039) at pH 5.4 and 6.7, 1 M Tris pH 8 (Thermo Fisher, AM9849), and 1 M Tris pH 9 (Millipore, 9295-OP) with stock solutions of 0.5 M ammonium iron(II) sulfate hexahydrate (Sigma-Aldrich, 09719), 0.5 M manganese(II) chloride tetrahydrate (Sigma-Aldrich, 63535), or 0.5 M magnesium chloride (Thermo Fisher, AM9530G). A typical RNA incubation consisted of 250 mM buffer, 1 mM EDTA (Thermo Fisher, AM9260G), 0 – 50 mM of metals, and 5 µM of the RNA oligomer in a final 20 µL reaction. All stock solutions were sparged with argon and stored inside an anaerobic glove box (Coy). The atmosphere inside the glove box was N_2_ with 2.5 – 3% H_2_ and internal circulation through a platinum catalyst maintained residual oxygen levels below 10 ppm. All RNA reactions occurred inside the glove box on a miniPCR mini16 thermocycler (Amplyus, QP-1016-01) kept at 75 °C in order to facilitate rapid RNA degradation.

### 2.5. RNA degradation quantification

All RNA degradation experiments were quantified via urea polyacrylamide gel electrophoresis (National Diagnostics, EC-830 & EC-840) followed by imaging on a Typhoon 9410 (GE Healthcare). Two bands were detected representing either the intact 15-mer (5’-Cy3-TTT-TTT-rCTT-TTT-TTT-3’) or the residual 7-mer (5’-Cy3-TTT-TTT-rC-3’) cleaved at the single ribonucleotide site (Figure 1). Scans were then analyzed with the ImageQuant TL 7.1 software (GE Healthcare).

### 2.5. RNA degradation kinetics

5 µM of the RNA oligomer was incubated in each of the Mars solutions as described above at 75 °C (n = 2) inside an anaerobic chamber. 1 µL aliquot time points were taken at 5, 15, 30, 60, and 120 minutes for pH 6.7, 8, and 9 and at 10, 30, 60, 120, and 240 minutes for pH 3.2 and 5.4. Aliquots were added to 25 µL of a kill buffer solution of 8 M Urea, 1x TBE, and 100 mM EDTA. Samples were then removed from the anaerobic chamber and 5 µL of the kill buffer and time point mixture (2 picomols RNA) was taken for gel electrophoresis and quantification. Pseudo-first order reaction kinetics *k*_obs_(h^−1^) of RNA hydrolysis were determined with the following relationship, ln[p_t_] = -*k*t + ln[p_o_] where p_t_ = percent intact oligo at t = time n and p_o_ = percent intact oligo at t = 0 (Figure 2).

**Figure 2:**
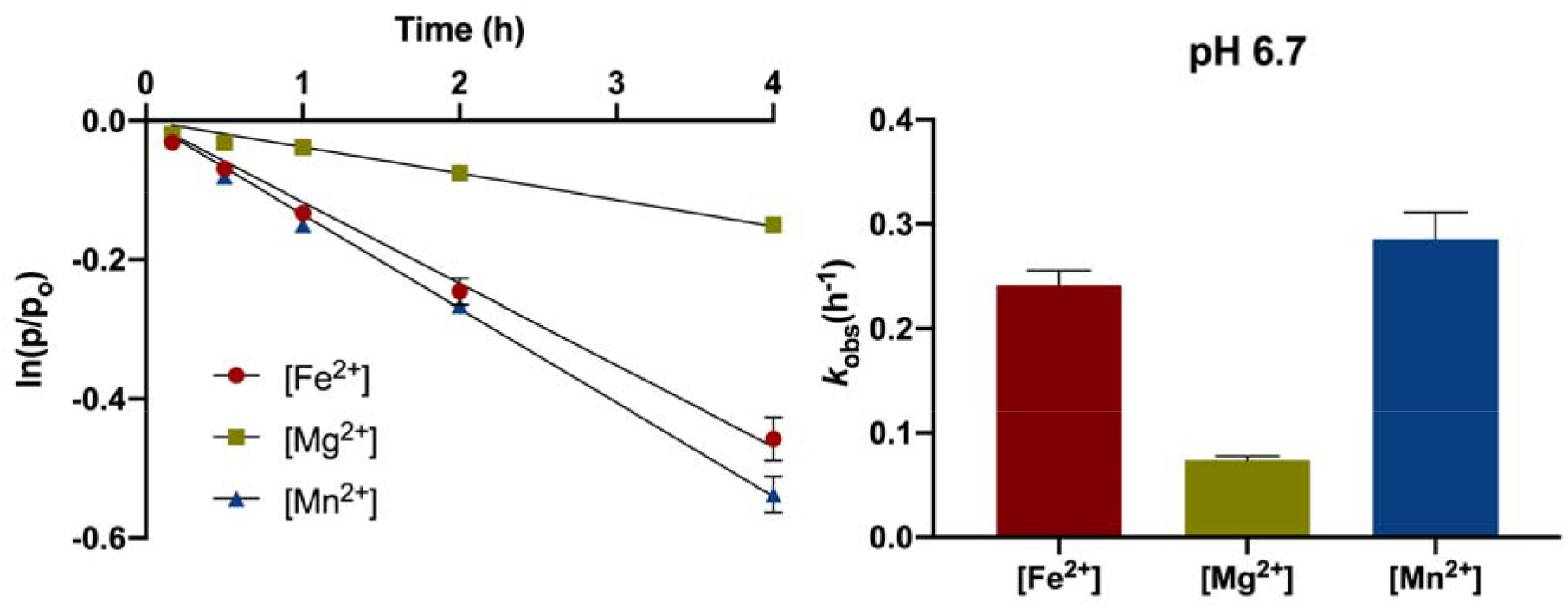
Metal ion catalysis of RNA degradation. Plot of oligonucleotide strand cleavage at the site of an RNA nucleotide as measured by urea polyacrylamide gel electrophoresis with 50 mM Fe^2+^ (•), 50 mM Mg ^2+^ (▴), and 50 mM Mn^2+^ (▪) at pH 6.7. The natural logarithm of the fraction of un-cleaved RNA with time (h) was fit to a linear regression ln[p_t_] = -*k*t + ln[p_o_] and the slope yielded our pseudo-first-order rate constants (*k*_obs_(h^−1^).

## 3. Results

Our experimental results indicate that enhanced RNA hydrolysis occurs due to the presence of metals (i.e., metal-catalyzed hydrolysis) in nearly all pH conditions (Table 1). However, at pH 3.2, increasing metal concentration decreases the rate of degradation (Figure 3). Hydrolysis at pH 3.2 is best mitigated by Mg^2+^ at 50 mM (*k*_obs_(h^−1^) = 0.013) followed by Fe^2+^ at 50 mM (*k*_obs_(h^−1^) = 0.025) then Mn^2+^ at 50 mM (*k*_obs_(h^−1^) = 0.026) relative to the negative control (*k*_obs_(h^−1^) = 0.081). Between pH 5.4 and 8, RNA incubations containing Mg^2+^ are generally more stable than those containing Fe^2+^. This preference is most apparent in saturation curves fitted to the Michaelis-Menten model for enzyme kinetics (Figure 4) which indicate a stability optimum at pH 5.4 where the maximum rate of RNA hydrolysis in the presence of Mg^2+^ (*k*_max_(h^−1^) = 0.0095) is notably lower than for Fe^2+^ (*k*_max_(h^−1^) = 0.071). Trends for Mn^2+^ solutions are quantitatively similar to Fe^2+^ between pH 3.2 and 8 (Figure 5, Table 1). At pH 9, we record the most rapid metal-catalyzed hydrolysis rates in our experiments. Michaelis-Menten models show that Fe^2+^ (*k*_max_(h^−1^) = 0.61) is more stable than Mg^2+^ (*k*_max_(h^−1^) = 0.96) although both Fe^2+^ and Mn^2+^ precipitated out of solution (Figure 4, Table 2). Results for the basalt analog solutions tend towards less overall metal-catalyzed hydrolysis in Mg^2+^-rich solutions (e.g., forsteritic) over Fe^2+^-rich solutions (e.g., fayalitic) (Figure 6). A summary of all the results produced in this study is presented in Table 1 and Table 2.

**Table 1.**

| Table 1 | pH 3.2 |  |  | pH 5.4 |  |  | pH 6.7 |  |  | pH 8 |  |  | pH 9 |  |  |
| --- | --- | --- | --- | --- | --- | --- | --- | --- | --- | --- | --- | --- | --- | --- | --- |
| Metal | $k_{\text{obs}} (\text{h}^{-1})$<br>(10<sup>-2</sup>) | S.E.M.<br>(10<sup>-3</sup>) | t<sub>1/2</sub><br>(h) | $k_{\text{obs}} (\text{h}^{-1})$<br>(10<sup>-2</sup>) | S.E.M.<br>(10<sup>-3</sup>) | t<sub>1/2</sub><br>(h) | $k_{\text{obs}} (\text{h}^{-1})$<br>(10<sup>-2</sup>) | S.E.M.<br>(10<sup>-3</sup>) | t<sub>1/2</sub><br>(h) | $k_{\text{obs}} (\text{h}^{-1})$<br>(10<sup>-2</sup>) | S.E.M.<br>(10<sup>-3</sup>) | t<sub>1/2</sub><br>(h) | $k_{\text{obs}} (\text{h}^{-1})$<br>(10<sup>-2</sup>) | S.E.M.<br>(10<sup>-3</sup>) | t<sub>1/2</sub><br>(h) |
| [Fe<sup>2+</sup>] |  |  |  |  |  |  |  |  |  |  |  |  |  |  |  |
| 50 mM | 2.47 | 1.40 | 28 | 5.07 | 1.70 | 14 | 24.1 | 14.5 | 3 | 30.0 | 6.10 | 2 | 28.9 | 13.2 | 2 |
| 25 mM | 3.41 | 0.70 | 20 | 4.26 | 0.50 | 16 | 16.5 | 1.30 | 4 | 27.7 | 12.6 | 3 | 13.3 | 2.65 | 5 |
| 10 mM | 4.49 | 3.05 | 15 | 2.58 | 1.85 | 27 | 4.40 | 3.65 | 16 | 14.1 | 13.8 | 5 | 12.1 | 5.50 | 6 |
| 5 mM | 4.24 | 2.55 | 16 | 1.44 | 0.95 | 48 | 2.26 | 2.75 | 31 | 6.76 | 11.3 | 10 | 7.48 | 1.20 | 9 |
| 2.5 mM | 4.17 | 1.10 | 17 | 0.54 | 0.25 | 130 | 0.74 | 3.80 | 94 | 0.65 | 0.25 | 107 | 5.40 | 7.00 | 13 |
| [Mg<sup>2+</sup>] |  |  |  |  |  |  |  |  |  |  |  |  |  |  |  |
| 50 mM | 1.28 | 0.50 | 54 | 0.75 | 0.70 | 92 | 7.42 | 3.75 | 9 | 6.76 | 0.30 | 10 | 62.3 | 11.3 | 1 |
| 25 mM | 1.97 | 0.65 | 35 | 0.92 | 0.05 | 76 | 5.61 | 1.45 | 12 | 4.41 | 3.00 | 16 | 44.9 | 25.1 | 2 |
| 10 mM | 2.43 | 2.05 | 29 | 0.73 | 0.10 | 95 | 3.31 | 1.30 | 21 | 2.16 | 0.70 | 32 | 29.0 | 31.2 | 2 |
| 5 mM | 2.77 | 0.40 | 25 | 0.63 | 0.40 | 110 | 2.46 | 1.45 | 28 | 1.85 | 6.10 | 37 | 13.4 | 13.9 | 5 |
| 2.5 mM | 3.40 | 2.15 | 20 | 0.20 | 0.10 | 347 | 1.44 | 3.20 | 48 | 1.06 | 4.10 | 65 | 5.32 | 0.60 | 13 |
| [Mn<sup>2+</sup>] |  |  |  |  |  |  |  |  |  |  |  |  |  |  |  |
| 50 mM | 2.57 | 0.05 | 27 | 5.22 | 0.35 | 13 | 28.6 | 25.5 | 2 | 82.7 | 20.7 | 1 | 87.5 | 29.9 | 1 |
| 25 mM | 2.03 | 1.70 | 34 | 5.68 | 0.80 | 12 | 12.2 | 24.1 | 6 | 58.9 | 9.05 | 1 | 127 | 62.1 | 1 |
| 10 mM | 4.24 | 2.00 | 16 | 2.64 | 2.90 | 26 | 6.29 | 18.0 | 11 | 25.4 | 3.10 | 3 | 104 | 27.9 | 1 |
| 5 mM | 4.22 | 2.28 | 16 | 2.07 | 3.10 | 33 | 2.95 | 6.95 | 24 | 14.1 | 11.0 | 5 | 92.2 | 4.65 | 1 |
| 2.5 mM | 7.03 | 3.10 | 10 | 0.53 | 1.10 | 131 | 0.72 | 0.25 | 97 | 4.20 | 3.00 | 17 | 6.04 | 12.9 | 11 |
| Basalt Analogs<br>[Mg<sup>2+</sup>:Fe<sup>2+</sup>] |  |  |  |  |  |  |  |  |  |  |  |  |  |  |  |
| 80:20 | 2.05 | 0.65 | 34 | 0.78 | 0.20 | 89 | 2.35 | 1.50 | 29 | 8.20 | 3.70 | 8 | 19.9 | 30.3 | 3 |
| 50:50 | 2.68 | 0.35 | 26 | 2.29 | 3.95 | 30 | 11.1 | 1.75 | 6 | 20.4 | 6.05 | 3 | 22.8 | 25.4 | 3 |
| 20:80 | 2.20 | 3.30 | 32 | 2.70 | 2.40 | 26 | 20.3 | 3.85 | 3 | 17.7 | 9.30 | 4 | 19.7 | 11.2 | 4 |
| Negative Control |  |  |  |  |  |  |  |  |  |  |  |  |  |  |  |
| 0 mM | 8.06 | 4.85 | 9 | 0.16 | 0.70 | 433 | 0.20 | 0.15 | 355 | 1.82 | 1.30 | 38 | 0.81 | 1.10 | 86 |

**Table 2.**

| **Metal** | **pH** | **$k_{\max}$ ($\text{h}^{-1}$) ($10^{-2}$)** | **S.E.M. ($10^{-3}$)** | **Km** | **S.E.M.** |
| --- | --- | --- | --- | --- | --- |
| **[Fe<sup>2+</sup>]** | 5.4 | 7.14 | 4.46 | 18.8 | 2.74 |
|  | 6.7 | 90.9 | 469 | 134 | 89.7 |
|  | 8 | 47.3 | 73.3 | 24.1 | 8.03 |
|  | 9 | 47.4 | 147 | 38.7 | 22.1 |
| **[Mg<sup>2+</sup>]** | 5.4 | 0.95 | 0.99 | 3.97 | 1.59 |
|  | 6.7 | 9.86 | 6.47 | 17.7 | 2.77 |
|  | 8 | 11.7 | 21.1 | 37.8 | 12.7 |
|  | 9 | 96.1 | 86.2 | 27.3 | 5.06 |

**Figure 3:**
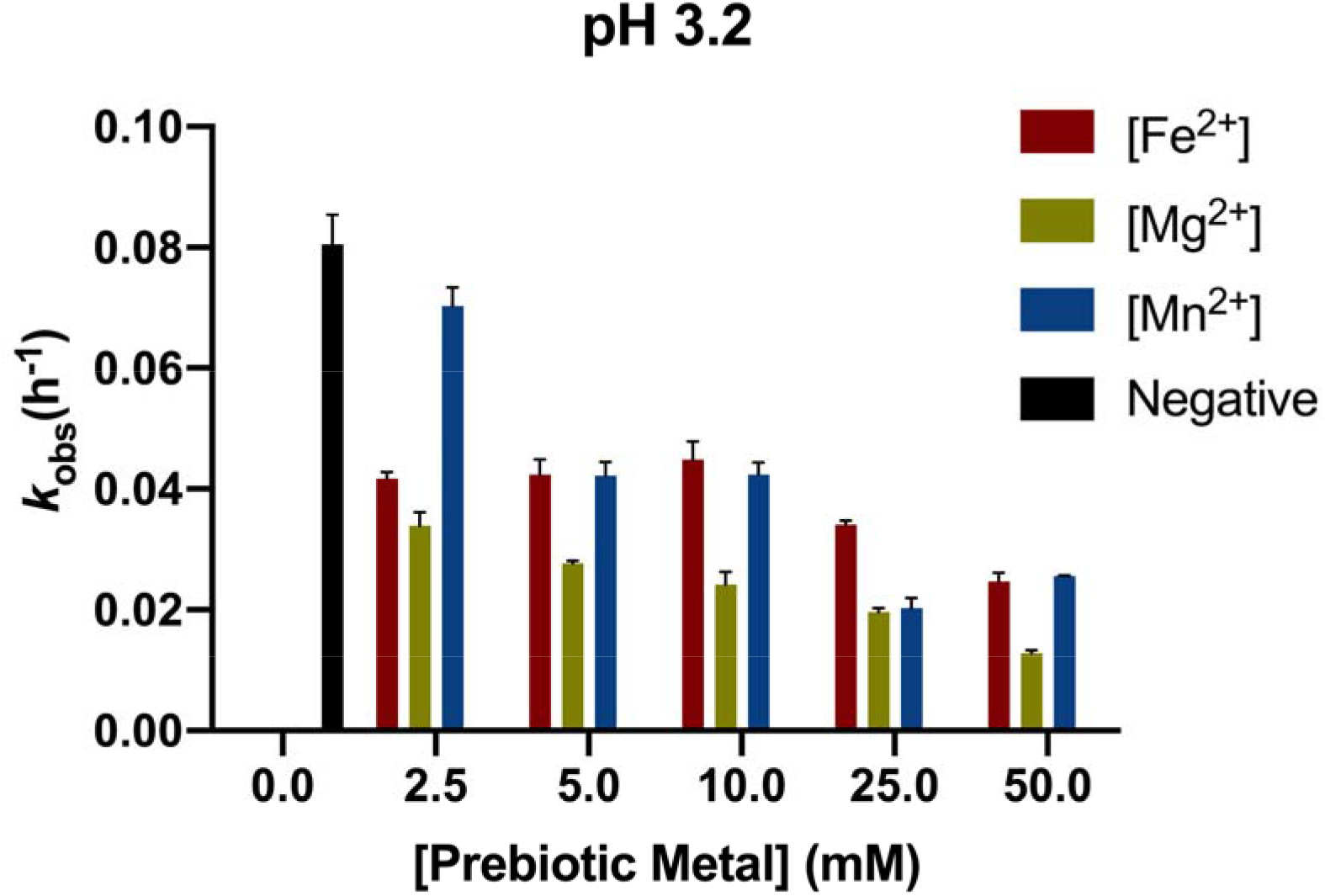
RNA degradation at pH 3.2. Incubation of the chimeric RNA-DNA oligonucleotide at pH 3.2 indicates that increasing concentrations of prebiotic metals decrease the rate of RNA degradation. RNA degradation is best mitigated by Mg^2+^ at 50 mM (*k*_*obs*_ = 0.013), compared to Fe^2+^ at 50 mM (*k*_*obs*_ = 0.025), Mn^2+^ at 50 mM (*k*_*obs*_ = 0.026) and the no divalent metal cation negative control (*k*_*obs*_ = 0.081). Error bars represent S.E.M. (n = 2).

**Figure 4:**
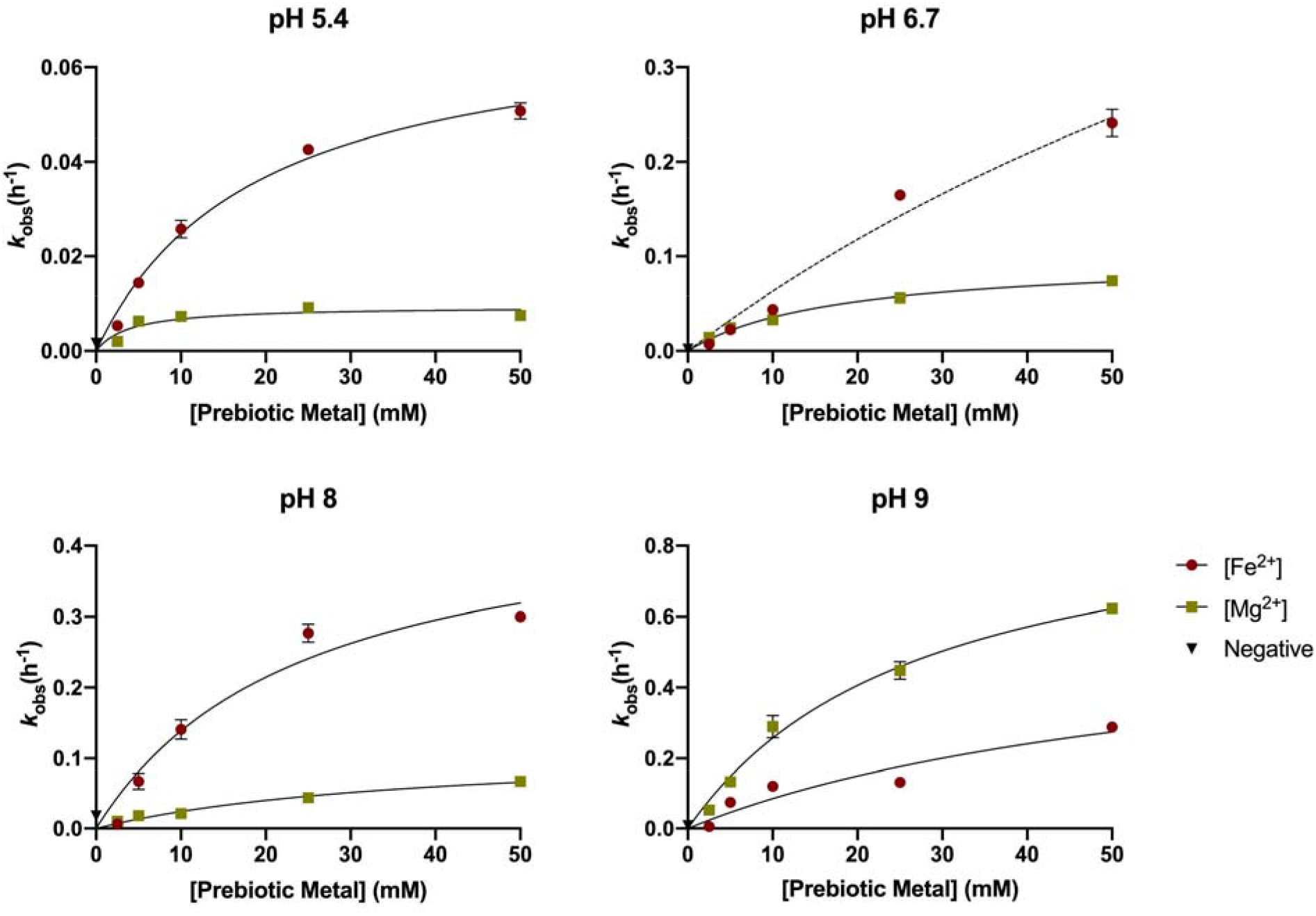
Characterization of RNA degradation kinetics. Saturation curves of Fe^2+^ and Mg^2+^ metal-catalyzed hydrolysis fitted to the Michaelis-Menten equation for enzyme kinetics. Within the range tested, we observe greater RNA stability at pH 5.4 where the maximum rate of hydrolysis (*k*_max_) with Mg^2+^ is lower than with Fe^2+^. RNA degradation experiments with Fe^2+^ at pH 6.7 did not reach saturation. Error bars represent S.E.M. (n=2).

**Figure 5:**
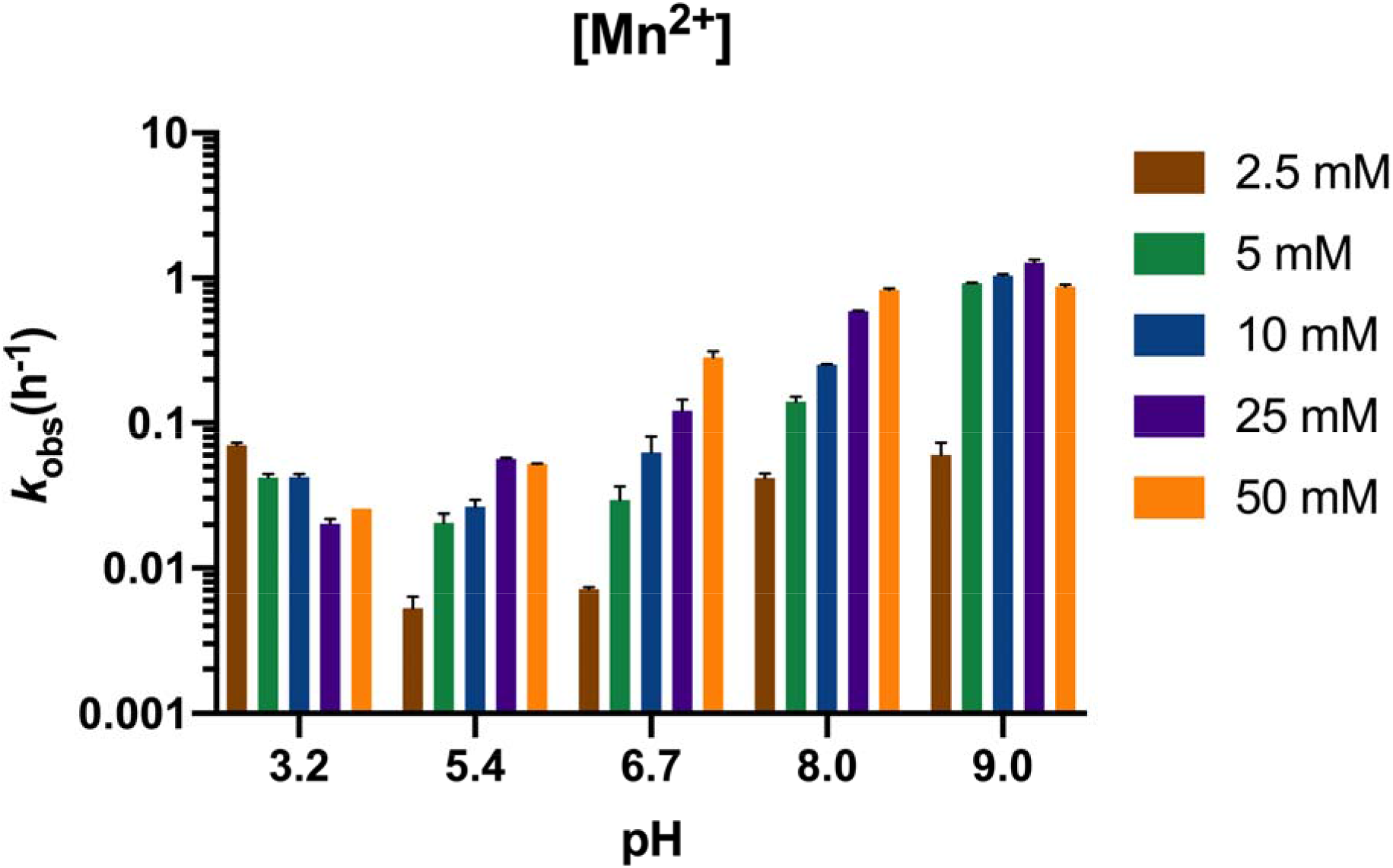
Manganese-catalyzed RNA degradation. Trends for Mn^2+^ are qualitatively similar to Fe^2+^ between pH 3.2 and 8. At pH 9 however, substantially more Fe^2+^ was observed to precipitate out of solution versus Mn^2+^. Error bars represent S.E.M. (n=2).

**Figure 6:**
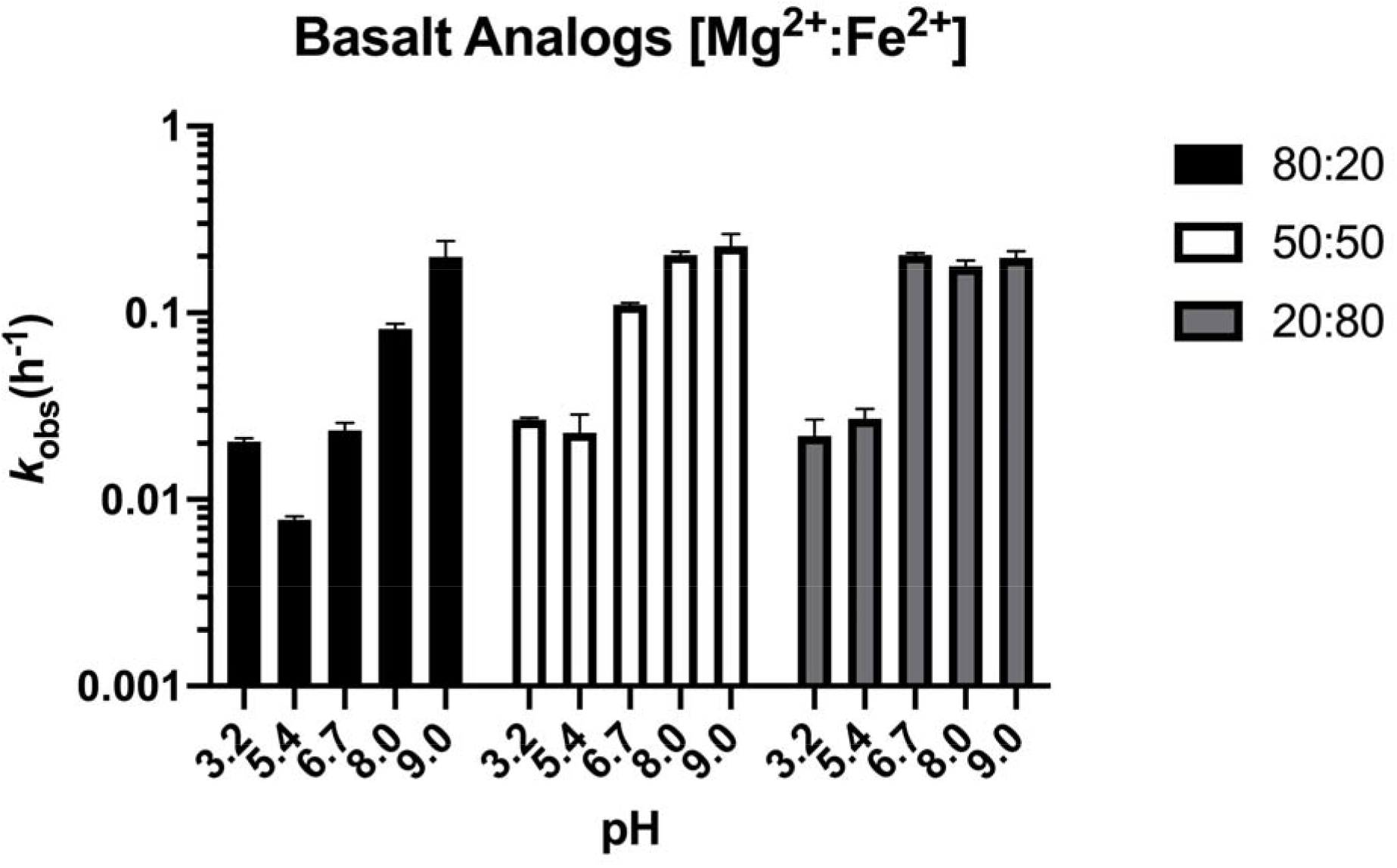
Prebiotic metal mixture-catalyzed RNA degradation. Results for the basalt analog solutions tend towards less overall metal-catalyzed hydrolysis in Mg^2+^-rich solutions (e.g., forsteritic) over Fe^2+^-rich solutions (e.g., fayalitic). Metal-catalyzed degradation at pH 9 for all basalt analogs is inferred to be primarily Mg^2+^-dominated as Fe^2+^ was observed to precipitate out of solution after 24 hours. Error bars represent S.E.M. (n=2).

## 4. Discussion

### 4.1. Prebiotic metal catalysis

In aqueous solution Mg^2+^, Fe^2+^, and Mn^2+^ form a hydrated hexa aquo species (Mg^2+^(H_2_O)_6_, p*K*_a_ = 11.4, Fe^2+^(H_2_O)_6_, p*K*_a_ = 9.6, and Mn^2+^(H_2_O)_6_, p*K*_a_ = 10.6) which polarize first shell water molecules in a tightly packed octahedral geometry (Jackson et al., 2015). Normally, these metal aquo complexes interact with secondary and tertiary RNA structures to neutralize the electrostatic repulsion of negatively charged phosphate groups brought into close proximity, or to increase local rigidity and join distal RNA structures by incorporating phosphate groups into their first coordination shell (Petrov et al., 2012). However, in our experiments, we sought to quantify the effect of metal-catalyzed RNA cleavage by transesterification, which is thought to be analogous to how certain ribozymes (e.g., the hammerhead self-cleaving ribozyme) utilize metals to stabilize transition states and catalyze cleavage of the 5’,3’-phosphodiester bond (Fedor, 2002; Hampel and Cowan, 1997; Johnson-Buck et al., 2011). Namely, 1) acid/base interactions with water result in the activation of the ribose 2’-hydroxyl nucleophile, and 2) first shell phosphate ligands draw electron density and expose phosphorous to nucleophilic attack. Attack of the 2’-hydroxyl on the adjacent phosphate results in formation of a 2’-3’ cyclic phosphate terminated oligonucleotide plus a second oligonucleotide product that begins with a 5’-hydroxyl.

Our results reproduce the enhanced RNA degradation rates expected to be associated with each metal’s respective acid dissociation constant (p*K*_a_) in solution. Between pH 5.4 and 8, slower rates of metal-catalyzed hydrolysis occur in the presence of Mg^2+^ (p*K*_a_ = 11.4) followed interchangeably by Mn^2+^ (p*K*_a_ = 10.6) and Fe^2+^ (p*K*_a_ = 9.6) (Figure 4, Table 1). Moreover, because of low lying d orbitals, both Fe^2+^ and Mn^2+^ have greater electron withdrawing power (Fe^2+^ = 0.11 e^−^) than Mg^2+^ (0.08 e^−^) (Okafor et al., 2017) which likely compound the rate of RNA cleavage due to mechanism 2) described above (Hampel and Cowan, 1997). At pH 9, it would appear that slower degradation rates occur in the presence of Fe^2+^ (*k*_max_(h^−1^) = 0.61) rather than Mg^2+^ (*k*_max_(h^−1^) = 0.96) contrary to our interpretation for pH 5.4 – 8 (Figure 4). Nonetheless, it is known that alkaline Fe^2+^ solutions will begin to form insoluble species such as Fe(OH)_2_ around pH ∼9 (Gayer and Woontner, 1956) that would sequester Fe^2+^ from participating in catalysis. Precipitates were observed for both Fe^2+^ and Mn^2+^ solutions albeit considerably more with Fe^2+^ at pH 9 corroborating interpretations that solubility properties are responsible for the observed differences between Mg^2+^ and Fe^2+^(Jin et al., 2018). Results for pH 3.2 were unexpected as increasing concentrations of prebiotic metals decreased the rate of hydrolysis (Figure 3) and additional work is required to understand the precise preservation mechanism. However, researchers have proposed that perhaps metal ions can act as Lewis acids which stabilize the 2’-hydroxyl group and prevent nucleophilic attack of phosphorous (Fedor, 2002). Results for the basalt analogs demonstrate greater degradation rates with increasing concentrations of Fe^2+^ relative to Mg^2+^ (Figure 6). This is best observed at pH 6.7 and 8 where 80:20 (20 mM Mg^2+^: 5 mM Fe^2+^) is mostly Mg^2+^-dominated then transitions to Fe^2+^-dominated at 20:80 (5 mM Mg^2+^: 20 mM Fe^2+^) (Table 1, Figure 6). Metal-catalyzed RNA cleavage at pH 9 for all basalt analogs is inferred to be primarily Mg^2+^-dominated as Fe^2+^ was observed to precipitate out of solution.

### 4.2. RNA on Mars

Our results demonstrate that RNA stability depends on both metal concentration and pH. While concentrations up to 50 mM were tested here, lower metal concentrations ∼1 mM are more geochemically plausible on Earth though estimates on Mars are not well constrained (Catling, 1999). Observations from anoxic crater lakes and perennially stratified ferruginous lakes on Earth show that ranges between 0 – 1.5 mM of dissolved Fe^2+^, Mn^2+^, or Mg^2+^ are reasonable for a basalt hosted basin (e.g., Bura-Nakic et al., 2009; Busigny et al., 2014; Hongve, 1994; Kling et al., 1989). However, such low values are often at odds with proposed prebiotic chemistries which require 50 - 250 mM of metals for *in situ* RNA synthesis (e.g., 250 mM Fe^2+^ - Patel et al., 2015) or replication (e.g., 50 - 200 mM Mg^2+^ - Szostak, 2012). It is unclear whether such high concentrations of metals would be geochemically reasonable on early Mars, however, periods of wet-dry cycling (e.g., playa environments) or endorheic lakes could conceivably facilitate required concentrations (e.g., Patel et al., 2015). Meanwhile, *in situ* observations on Mars suggest pH regimes ranging from acidic (pH 2 - 4) (e.g., Squyres and Knoll, 2005) to circumneutral (pH ∼7) (e.g., Grotzinger et al., 2014,) existed on various locations around the planet. Observations by MAVEN constrain the composition of the early atmosphere to primarily CO_2_-dominated at ≥0.5 bars, which accordingly would not considerably acidify surface environments below pH ∼6.7 (Kua and Bada, 2011). Modeling of continental weathering of early Earth basalts additionally suggests that waters with high alkalinity would have stabilized pH between 6.6 and 7 under a ∼1 bar CO_2_ atmosphere (Halevy and Bachan, 2017; Krissansen-Totton et al., 2018). This is important because subsurface basaltic aquifers on Mars would have globally theoretically sustained neutral pH waters as observed at Gale Crater (Grotzinger et al., 2014).

Our results indicate that a Mg^2+^-rich basalt (e.g., McSween, 2002; Mustard et al., 2005) sourcing metals to a slightly acidic (pH 5.4) aqueous environment on Mars would have best supported long-lived single-stranded RNA polymers. Notwithstanding this conclusion, CO_2_ pressures (10 bar) required to acidify surface waters are not supported by atmospheric models (Forget et al., 2013; Tian et al., 2010) while buffering from a basaltic aquifer would neutralize pH as indicated above. Results from our experiments at pH 6.7 therefore represent the most accurate interpretation of potentially global conditions on Mars (Bibring et al., 2006), or at least those found locally at Gale Crater (Hurowitz et al., 2017). Depending on how early Mars became oxidizing, aqueous environments like Meridiani Planum and Gusev Crater (pH 2 - 4) would accelerate RNA hydrolysis rates at low concentrations as observed at pH 3.2 (Figure 3) while the oxidation of Fe^2+^ to Fe^3+^ would produce a well-known iron species that forms a strong complex with phosphate and leads to the precipitation of RNA (van Roode and Orgel, 1980). This is particularly significant because researchers have proposed that Fe^2+^ may have promoted RNA replication (Jin et al., 2018), folding (Athavale et al., 2012), and novel catalytic activity (Hsiao et al., 2013; Okafor et al., 2017) on Earth prior to the great oxidation event ∼2.45 Gya when life would have transitioned to Mg^2+^.

Most relevant to this study, work on non-enzymatic RNA replication has demonstrated that metal-catalyzed hydrolysis increases the rate of polymerization by facilitating the deprotonation of the 3’-hydroxyl group in 2-methylimidazole nucleotides (Li et al., 2017) activating the 3’-hydroxyl as a nucleophile. Work by Jin et at., 2018 has shown that Fe^2+^ facilitates RNA primer extension in solutions containing template strands and monomers at a weakly acidic to neutral (pH ∼7) optimum while Mg^2+^ is most effective at alkaline conditions (pH ∼9). Fe^2+^ could have accordingly enabled an RNA world (as described by Athavale et al., 2012) on Mars, though oxidizing conditions would have favored Mg^2+^ as a catalyst at alkaline conditions as early as ∼4.1 Gya. Moreover, for the RNA World to have existed, synthesis (i.e. replication) is at least, if not more important than stability. While degradation in general is slower at weakly acidic pH values, so is template copying chemistry (Jin et at., 2018). Replication with Mg^2+^ as the metal cofactor works best at mildly alkaline pH values, where degradation of RNA is still fairly slow. Fe^2+^ works best as a replication cofactor at neutral to very slightly acidic pH, however, it has severe effects on the stability of RNA and of activated monomers. In other words, replication must be faster than degradation.

Future work is required to further constrain the composition of theoretical Mars waters with respect to mechanisms that may have accumulated metals to prebiotically relevant concentrations (e.g., playas, brines). The work presented here highlights the importance of metals and pH derived from variable bedrock compositions and hypothetical atmospheric conditions on RNA stability. Additional studies will seek to include non-enzymatic RNA extension, the effect of template and complement RNA strands, and additional geological parameters such as UV flux. In summary, the work presented here advances our understanding of how geochemical environments could have influenced the stability of a potential RNA world on Mars.

## 5. Conclusions

Discoveries of structural and regulatory RNA molecules suggests that life as we know it may have emerged from an earlier RNA world (Bernhardt, 2012). Subsequently, strides in prebiotic chemistry have demonstrated the synthesis and stability of, and catalysis by, RNA polymers under various conditions and in presence of cofactors (Bernhardt and Tate, 2012). However, it has been challenging to confidently determine the types of real-world environments that may have existed on the Hadean Earth and hosted the origin of life due to global resurfacing and recycling (Marchi et al., 2014). We believe that Mars is the next best alternative to search for environments consistent with requirements imposed by the RNA world. In this study we investigated the influence of bedrock composition (e.g., mafic-ultramafic, iron-rich, magnesium-rich, etc.) and subsequent weathering of prebiotically relevant metals (Fe^2+^, Mg^2+^, and Mn^2+^) on RNA stability. These metals have been demonstrated to catalyze hydrolysis, folding, and enable RNA catalytic activity. In addition, we simultaneously adjusted pH to reflect the composition of hypothetical CO_2_-dominated atmospheres in equilibrium with surface waters and an acidic and alkaline vent. We determined that metal-catalyzed hydrolysis of RNA depends on metal concentration and pH. Our results reproduce the enhanced RNA cleavage rates associated with each metal’s respective acid dissociation constant (p*K*_a_), and the increase in RNA degradation with increasing metal concentration (Figure 4, Table 1). Meanwhile, degradation rates decreased with increasing metal concentration via an unknown preservation mechanism at pH 3.2 (Figure 3). At pH 9, we encountered Fe^2+^ precipitation which artificially decreased hydrolysis rates while Mn^2+^ precipitation occurred to a lesser extent. We conclude that a Mg^2+^-rich basalt sourcing metals to slightly acidic (pH 5.4) waters would therefore be the stability optimum (as determined here) for RNA on Mars. However, it is important to note RNA replication chemistry with Mg^2+^ as the metal cofactor requires mildly alkaline pH values (Jin et at., 2018) in order to result in net accumulation. Fortunately, geologic evidence and modeling of basalt weathering indicate that Mars waters would have been near neutral pH ∼7. Our experiments at pH 6.7 therefore represent the most accurate interpretation of potentially global conditions on Mars. Results from Fe^2+^ at this pH and prior work on iron catalysis suggest that while high hydrolysis rates lower RNA stability, the presence of Fe^2+^ could have imparted novel catalytic function to candidate ribozymes. However, global oxidizing conditions (due to the lack of a dynamo) on the surface of Mars may have led to significant RNA instability due to the precipitation of RNA-Fe^3+^ complexes in Fe^2+^-rich environments as early as ∼4.1 Gya. We therefore presume the non-redox-sensitive Mg^2+^ would have been a principal catalyst similar to as on the Earth after the great oxidation event. Future Mars exploration should seek to determine the existence and distribution of such predicted environments in order to assess the possible nature and abundance of conditions conducive to an RNA world.

## Acknowledgements

This work was supported by NASA MatISSE award NNX15AF85G and the MIT Dean of Science Fellowship. We thank Christopher E. Carr and Shui-Ying (Fanny) Ng for helpful input.

